# Cortical patterning of abnormal morphometric similarity in psychosis is associated with brain expression of schizophrenia related genes

**DOI:** 10.1101/501494

**Authors:** Sarah E Morgan, Jakob Seidlitz, Kirstie Whitaker, Rafael Romero-Garcia, Nicholas E Clifton, Cristina Scarpazza, Therese van Amelsvoort, Machteld Marcelis, Jim van Os, Gary Donohoe, David Mothersill, Aiden Corvin, Andrew Pocklington, Armin Raznahan, Philip McGuire, The PSYSCAN Consortium, Petra E Vértes, Edward T Bullmore

## Abstract

Schizophrenia has been conceived as a disorder of brain connectivity but it is unclear how this network phenotype is related to the emerging genetics. We used morphometric similarity analysis of magnetic resonance imaging (MRI) data as a marker of inter-areal cortical connectivity in three prior case-control studies of psychosis: in total, N=185 cases and N=227 controls. Psychosis was associated with globally reduced morphometric similarity (MS) in all 3 studies. There was also a replicable pattern of case-control differences in regional MS which was significantly reduced in patients in frontal and temporal cortical areas, but increased in parietal cortex. Using prior brain-wide gene expression data, we found that the cortical map of case-control differences in MS was spatially correlated with cortical expression of a weighted combination of genes enriched for neurobiologically relevant ontology terms and pathways. In addition, genes that were normally over-expressed in cortical areas with reduced MS were significantly up-regulated in a prior post mortem study of schizophrenia. We propose that this combination of neuroimaging and transcriptional data provides new insight into how previously implicated genes and proteins, as well as a number of unreported proteins in their vicinity on the protein interaction network, may interact to drive structural brain network changes in schizophrenia.

## Introduction

Psychotic disorders have a lifetime prevalence of 1-3% and can be extremely debilitating (1). However, despite significant efforts, the brain architectural changes and biological mechanisms causing psychotic disorders are not yet well understood (2) and there has been correspondingly limited progress in the development of new therapeutics (3).

Magnetic resonance imaging (MRI) studies of schizophrenia have robustly demonstrated local structural differences in multiple cortical areas, subcortical nuclei and white matter tracts (4). The most parsimonious explanation of this distributed, multicentric pattern of structural change is that it reflects disruption or dysconnectivity of large-scale brain networks comprising anatomically connected brain areas. However, testing this dysconnectivity hypothesis of psychotic disorder has been constrained by the fundamental challenges in measuring anatomical connectivity and brain anatomical networks in humans. The principal imaging methods available for this purpose are tractographic analysis of diffusion weighted imaging (DWI) and structural covariance analysis of conventional MRI. DWI-based tractography generally under-estimates the strength of long distance anatomical connections, for example between bilateral homologous areas of cortex. Structural covariance analysis is not applicable to single subject analysis and its biological interpretation is controversial.

We recently proposed a technique known as “morphometric similarity mapping” (5), which quantifies the similarity between cortical areas in terms of multiple MRI parameters measured at each area and can be used to construct whole brain anatomical networks for individual subjects. In keeping with histological results indicating that cytoarchitectonically similar areas of cortex are more likely to be anatomically connected (6), morphometric similarity (MS) in the macaque cortex was correlated with tract-tracing measurements of axonal connectivity. Compared to both tractographic DWI-based networks and structural covariance networks, MS networks included a greater proportion of connections between human cortical areas of the same cytoarchitectonic class. Individual differences in the connectivity (or “hubness”) of cortical nodes in MS networks accounted for about 40% of the individual differences in IQ in a sample of 300 healthy young people. These results suggest that MS mapping could provide a useful new tool to analyse functionally relevant group-differences in brain structure.

Here we used MS mapping to test the dysconnectivity hypothesis of psychosis in three independent case-control MRI datasets: the Maastricht GROUP study (83 cases, 68 controls) and the Dublin study (33 cases and 82 controls), both made available as legacy datasets for the PSYSCAN project, and the publicly available Cobre dataset (69 cases and 77 controls); see Methods. We mapped case-control MS differences at global and nodal levels of resolution individually in each dataset to assess replicability and we tested for significant differences in network organization that were consistent across studies. We used partial least squares (PLS) regression to test the hypothesis that this MRI network phenotype of psychosis was correlated with anatomically patterned gene expression using data from the Allen Human Brain Atlas (AHBA). This approach was pioneered by (7, 8) and has already been applied in the context of disease (9, 10), despite the fact that the AHBA is based on healthy brains. Finally we tested the more pathogenically specific hypothesis that the genes most strongly associated with case-control differences in MS were enriched for genes that have been ontologically linked to neurobiological processes or genes that are abnormally expressed in postmortem studies of schizophrenia.

## Results

### Case-control differences in global morphometric similarity

Globally, MS was reduced in cases compared to controls in all 3 datasets (Fig. S2). Regional MS had an approximately Normal distribution over all 308 regions and in all 3 datasets there was a significant case-control difference in this distribution (P *<* 0.001, Kolmogorov-Smirnoff test). Modal values of regional MS were more frequent, and extreme values less frequent, in cases compared to controls (Fig. S2).

### Case-control differences in regional morphometric similarity

The cortical map of regional MS in Fig. 1 a) summarises the anatomical distribution of areas of positive and negative similarity on average over controls from all 3 datasets. The results are similar to those reported in an independent sample (5), with high positive MS in frontal and temporal cortical areas and high negative MS in occipital, somatosensory and motor cortex. This confirms the replicability of this pattern of regional MS in healthy individuals and is consistent with prior knowledge that primary cortex is more histologically differentiated than association cortex.

**Fig. 1.**
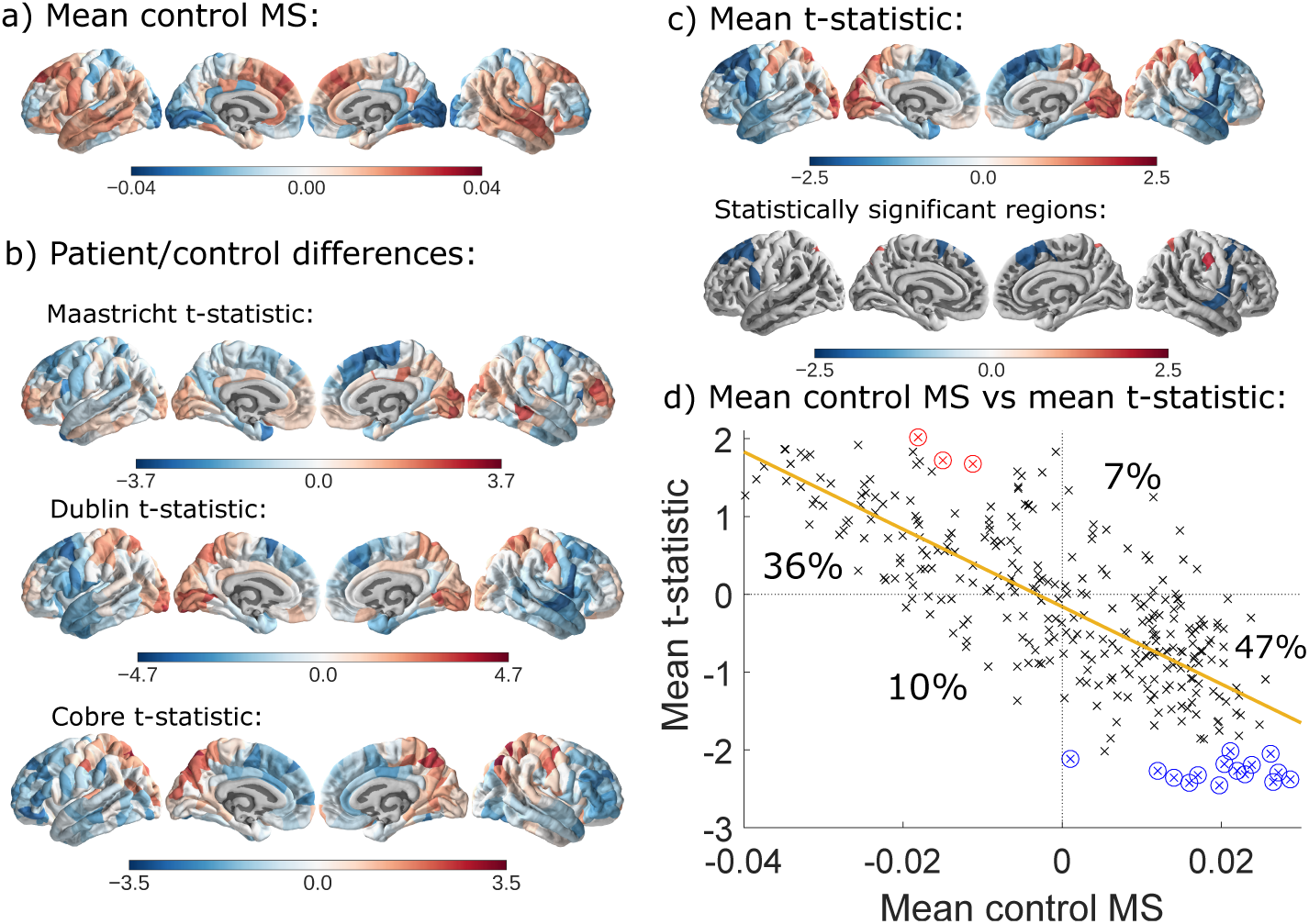
Case-control differences in regional MS. a) Regional MS averaged over healthy controls from all 3 datasets. b) *t*-statistics for the case-control difference in regional MS for each dataset. c) *t*-statistics for case-control difference in regional MS averaged over all 3 datasets and the *t*-statistics for case-control difference in regional MS in the 18 cortical areas where the difference was statistically significant on average over all 3 datasets (FDR = 0.05). d) Scatterplot of control mean regional MS (*x*-axis) versus case-control *t*-statistic (*y*-axis). Control MS (from panel a) is strongly negatively correlated with case-control differences in MS (from panel c) (Pearson’s *r*=-0.76, *P <* 0.001. Most cortical regions have positive MS in healthy controls and decreased MS in patients with psychosis (47% of regions) or negative MS in controls and increased MS in patients (36% of regions). Statistically significant regions are circled in red/blue according to whether their mean *t*-statistic increases/decreases in patients.

We mapped the i-statistics for the case-control differences in regional MS at each cortical area (Fig. 1 b). A positive i-statistic means MS increased in patients whereas a negative i-statistic means MS decreased. We found somewhat similar patterns of case-control difference across all 3 datasets, with increased regional MS in occipital and parietal areas in patients, and decreased regional MS in frontal and temporal cortex. The Dublin dataset showed the greatest number of cortical regions with significant case-control differences (Fig. S3). The convergence of case-control differences was quantified by correlating the *t*-statistics between datasets. The Dublin *t*-statistic was significantly correlated with both the Maastricht and the Cobre *t*-statistics (*r* = 0.42, *P <* 0.001 and *r =* 0.47, *P <* 0.001, respectively), although the Maastricht and Cobre *t*-statistics were not significantly correlated with each other (r = 0.058, *P* = 0.31), see Fig. S4. These correlations were robust to a permutation test designed to account for spatial autocorrelation amongst neighboring cortical areas (11), as well as a permutation test in which group labels were scrambled; see Supplemental Information (SI). We combined the *P*-values for case-control differences across all 3 datasets using Fisher’s method, and corrected for multiple comparisons using the false discovery rate (FDR), to identify 18 cortical regions where MS was robustly and significantly different between groups (Fig. 1 c). MS decreased in patients in 15 regions located in the superior frontal, caudal middle frontal, pre-central, pars triangularis and superior temporal areas and increased in 3 regions located in superior parietal and post-central areas (Table S2).

There was a strong negative correlation between regional MS in the control subjects and the case-control differences in regional MS (both averaged over all 3 datasets) (Fig. 1 d). Hence areas with highest positive MS in controls tended to show the greatest decrease of MS in patients; and, conversely, areas with highest negative MS in healthy controls had the greatest increase of MS in psychosis.

### Gene expression related to morphometric similarity

We used PLS regression to identify patterns of gene expression that were correlated with the anatomical distribution of case-control MS differences. The first PLS component explained 13% of the variance in the case-control MS differences, combining data from all 3 studies, significantly more than expected by chance (permutation test, *P <* 0.001). PLS1 gene expression weights were positively correlated with case-control MS differences in the Dublin study (*r* = 0.49, *P <* 0.001) and the Cobre study (*r* = 0.37, *P <* 0.001) (Fig. 2a); but not in the Maastricht study (*r* = 0.006, *P* = 0.94). These positive correlations mean that positively weighted genes are over-expressed in regions where MS increased in patients, and negatively weighted genes are over-expressed in regions where MS decreased in patients (Fig. 2d).

**Fig. 2.**
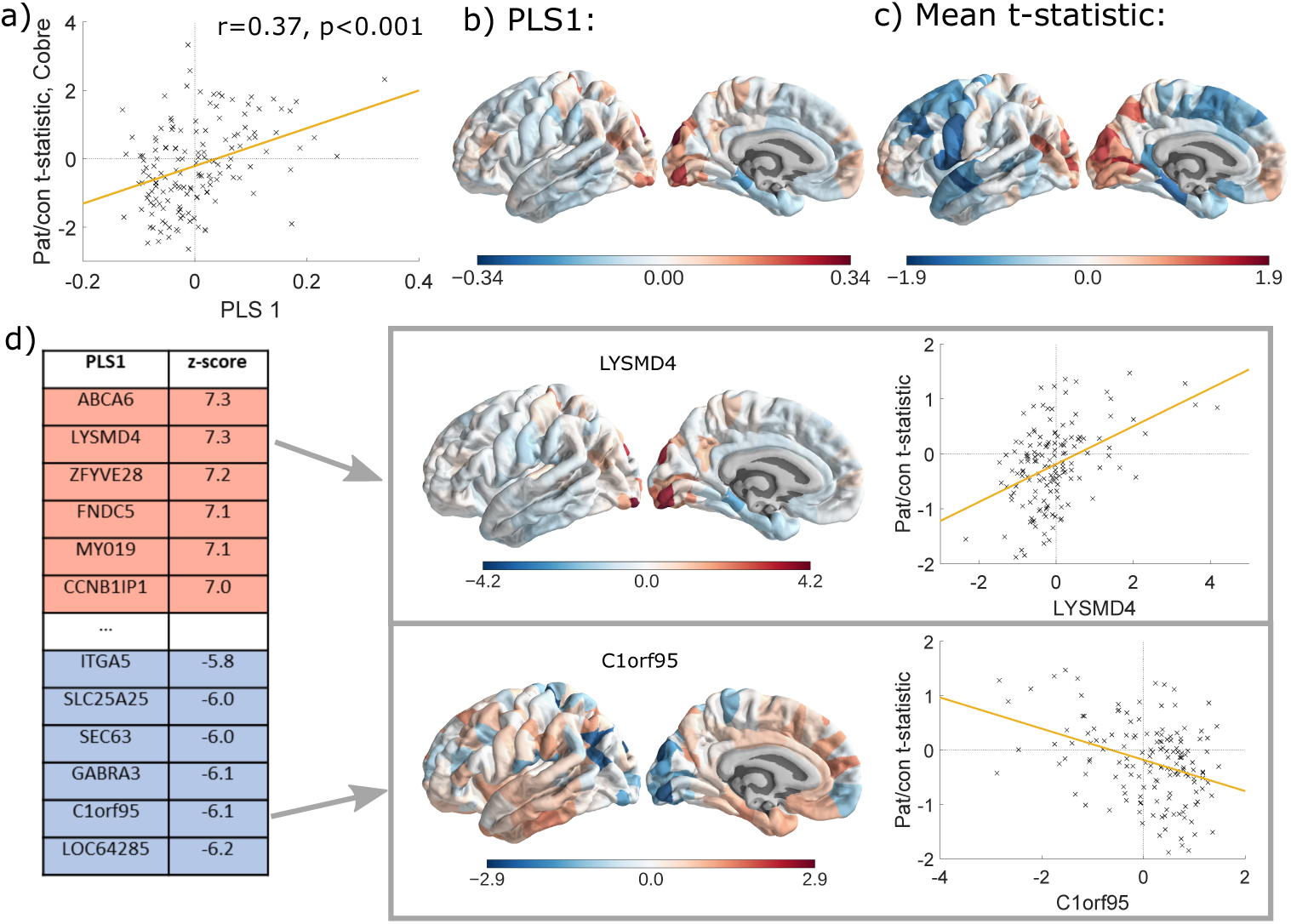
Gene expression profiles related to case-control differences in MS. a) Scatterplot of regional PLS1 scores (weighted sum of 20,647 gene expression scores) versus case-control differences in regional MS (Cobre dataset). b) Cortical map of regional PLS1 scores. c) Cortical map of mean control/patient MS differences (left hemisphere only, averaged across all datasets). d) Genes that are strongly positively weighted on PLS1 (e.g., *LYSMD4)* correlate positively with case-control differences in regional MS (*r =* 0.44, *P <* 0.001); whereas genes that are strongly negatively weighted on PLS1 (e.g., *C1orf95*) correlate negatively with case-control differences in MS (*r* = -0.37, *P <* 0.001).

### Enrichment analysis of genes transcriptionally related to morphometric similarity

We focused first on the genes that were most negatively weighted on PLS1, the PLS-gene set, because these are the genes that are normally highly expressed in areas where MS is reduced in patients. We found 1110 genes with normalised PLS1 weights *Z <* –3 and mapped the network of known interactions between proteins coded by this PLS-gene set (12) (Fig. 3). The resulting protein-protein interaction (PPI) network had 341 connected proteins and 1022 edges, significantly more than the 802 edges expected by chance (permutation test, *P < 1.0e –* 13). We also tested the PLS-gene set for significant GO enrichment of biological processes and enrichment of KEGG pathways, compared to a background of brain-expressed genes. Enriched biological processes included “nervous system development”, “synaptic signaling” and “adenylate cyclase-modulating G-protein coupled receptor signaling pathway” (see Dataset S1). There were two significantly enriched KEGG pathways: “neuroactive ligand-receptor interaction” and “retrograde endocannabinoid signaling” (Fig. S7). The proteins coded by genes enriched for “adenylate cyclase and related G-protein coupled receptor signalling pathways” and the two KEGG pathways formed the most strongly interconnected cluster of nodes in the PPI network, compatible with them sharing a specialised functional role for GPCR signaling.

**Fig. 3.**
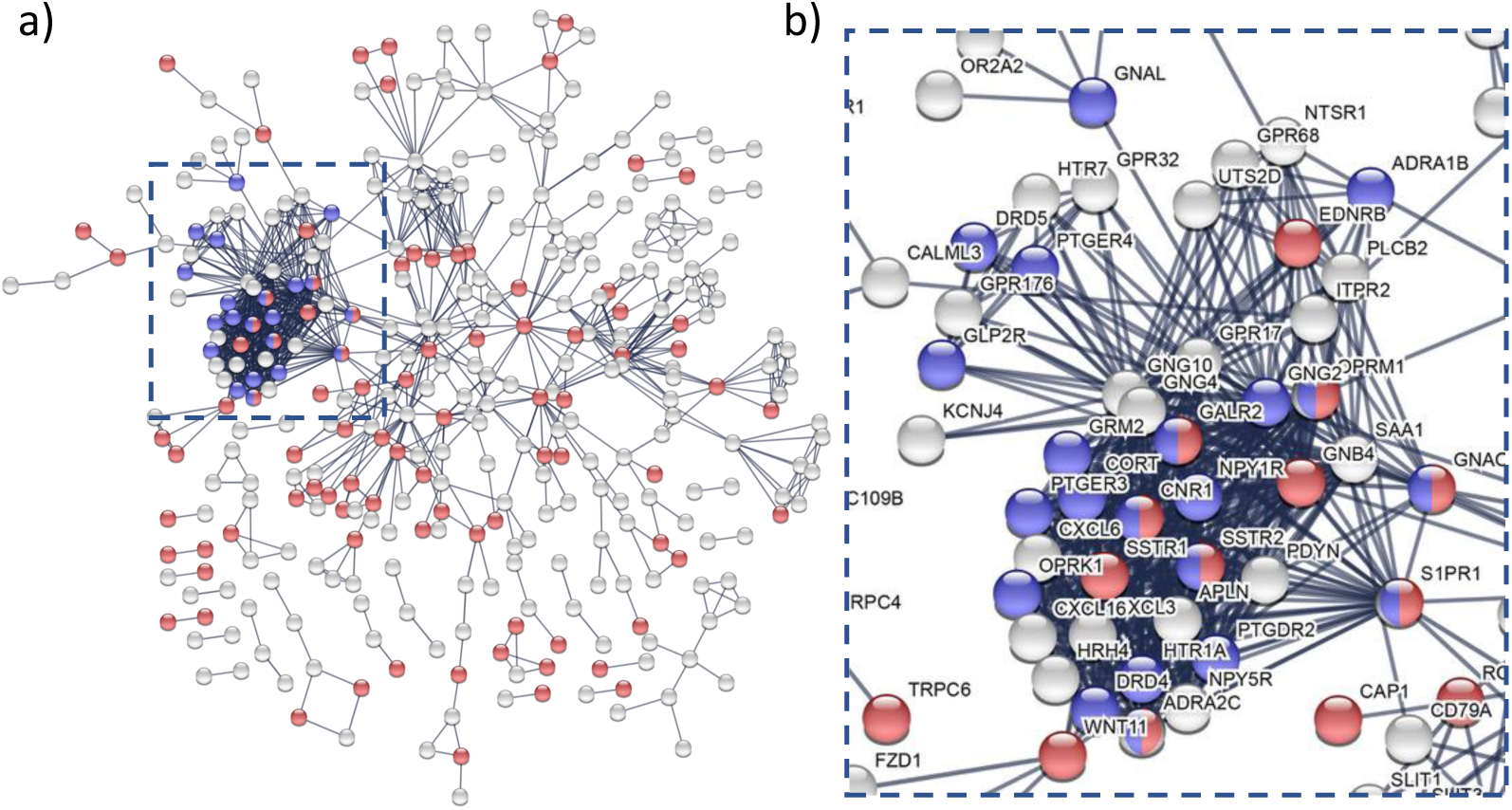
Enrichment analysis of genes transcriptionally related to MS. a) protein-protein interaction (PPI) network for PLS-genes (Z *<* -3), highlighted with some of the significantly GO enriched biological processes: “nervous system development” in red and “adenylate cyclase-modulating G-protein coupled receptor signaling pathway” in blue. b) The most interconnected set of proteins were coded by several genes previously implicated in schizophrenia: *PLCB2* (14, 16), *GRM2* (17), *DRD4* (18), *DRD5* (19), *ADRA2C* (20), *HTR1A* (21), *HTR7* (22), *CNR1* (23), *OPRM1* (24), *PT-GER3* (25), *NPY1R* (26), *SSTR2* (27), *NOS1* (28), *GNG2* (29), *PDYN* (30), *EDNRB* (13), *S1PR1* (13), *ITPR2* (13) and *NTSR1* (31).

To test more formally for enrichment of genes linked to schizophrenia, we used prior information about both gene transcription and sequence variation. Genes recently reported as over-expressed in post mortem brain tissue from patients with schizophrenia (13) were highly enriched among genes that were negatively weighted on PLS1 (permutation test, *P <* 10^−4^, before FDR correction), in comparison to a background of brain expressed genes (see SI for all gene expression results after FDR correction). In other words, genes that were up-regulated in post mortem brain tissue from patients with schizophrenia are normally over-expressed in association cortical areas that have reduced MS in psychosis. This result was not specific to schizophrenia and there is substantial overlap between genes which are up/down regulated in schizophrenia and other psychiatric disorders (e.g. autistic spectrum disorders, ASD), see the SI.

Genes which were over-expressed in regions where MS decreased in patients were not significantly enriched for common variant association with schizophrenia, derived from a recent genome-wide association study (GWAS) of PGC and CLOZUK samples (*P* > 0.05). However, schizophrenia risk genes from the DISEASES dataset (14), which combines results from gene association studies and text mining of biomedical abstracts, were significantly enriched among genes that were negatively weighted on PLS1 (permutation test, *P* = 0.048). There was no significant enrichment of risk genes for several other disorders in the DISEASES dataset (Alzheimer’s disease, ASD, attention deficit hyperactivity disorder, bipolar disorder), although risk genes for “mental depression” were significantly enriched (permutation test, *P* = 0.015); see SI.

We also investigated the PLS+ gene set that was normally over-expressed in parietal and other cortical areas where MS was abnormally increased in psychosis. The corresponding PPI network again showed significantly more interactions than expected by chance (P< 1.0e – 6), and was enriched for the biological process “nucleic acid metabolic process” but no KEGG pathways (see the SI). Genes which are down-regulated post mortem in schizophrenia (13) were highly enriched among genes that were positively weighted on PLS1 (permutation test, *P <* 0.001). Risk genes identified by the PGC and CLOZUK GWAS study (15), or by the DISEASES database, were not significantly enriched among these genes (*P* > 0.05).

## Discussion

### Morphometric similarity network phenotypes

Morphometric similarity mapping disclosed a robust and replicable cortical pattern of differences in psychosis patients. MS was significantly reduced in frontal and temporal cortical areas, and significantly increased in parietal cortical areas. This pattern was consistent across 3 independent datasets, with different samples, locations, scanners and scanning parameters; although it was least clearly evident in the dataset with the lowest image quality scores.

What does this novel MRI phenotype of psychosis represent? Morphometric similarity quantifies the correspondence or kinship of two cortical areas in terms of multiple macro-structural features, e.g., cortical thickness, and micro-structural features, e.g., fractional anisotropy (FA), that are measurable by MRI. We assume that high MS between a pair of cortical regions indicates that there is a high degree of correspondence between them in terms of cytoarchitectonic and myeloarchitectonic features that we cannot directly observe, given the limited spatial resolution and cellular specificity of MRI. Prior work also showed that morphometrically similar cortical regions are more likely to be axonally connected to each other, i.e., MS is a proxy marker for anatomical connectivity (5). Namely, histological evidence showed that cytoarchitectonically similar cortical areas are more likely to be axonally connected, and prior MRI data showed that MS is greater between human cortical areas belonging to the same cytoarchitectonic class, and between macaque cortical areas that are known to be axonally connected by tract-tracing experiments. We therefore interpret the reduced MS we observe in regions like the frontal and temporal nodes in psychosis as indicating that there is reduced architectonic similarity, or greater architectonic differentiation, between these areas and the rest of the cortex, which is probably indicative of reduced anatomical connectivity to and from the less similar, more differentiated cortical areas.

There is a well-evidenced and articulated prior theory of schizophrenia as a dysconnectivity syndrome, specifically functional dysconnectivity of frontal and temporal cortical areas has been recognised as a marker of brain network disorganization in schizophrenia. Our results of reduced MS in frontal and temporal cortex - implying increased architectonic differentiation and decreased axonal connectivity - are descriptively consistent with this theory. Our complementary finding of abnormally increased MS in parietal cortex - implying increased architectonic similarity and axonal connectivity - is plausible but not so clearly precedented, given the relatively limited prior data on the parietal cortex in studies of schizophrenia as a dysconnectivity syndrome (32, 33).

Encouragingly, this novel MRI network marker of psychosis was highly reliable across three independent and methodologically various case-control studies. This implies that the measurement is robust enough to be plausible as a candidate imaging biomarker of cortical network organization in large-scale, multi-centre studies of psychosis. However, this is not to say that quality of imaging data is not important. The MS “signal” of psychosis was less clearly detectable in the Maastricht GROUP study, which had the lowest scores on structural MRI data quality, and most clearly detectable in the Dublin study, which had the highest MRI quality scores.

### Transcriptional profiling of MS network phenotypes

In an effort to connect these descriptive results to the emerging genetics and functional genomics of schizophrenia, we first used PLS to identify the weighted combination of genes in the whole genome that has a cortical expression map most similar to the cortical map of case-control MS differences. Then we tested the mechanistic hypothesis that the genes with greatest (positive or negative) weight on PLS1 were enriched for genes previously implicated in the pathogenesis of schizophrenia.

We found that the genes that are normally over-expressed in frontal and temporal areas of reduced MS in psychosis, were significantly enriched for genes that are up-regulated in post mortem brain tissue from patients with schizophrenia (13). Conversely, the genes that are normally over-expressed in parietal and other areas of increased MS in psychosis were significantly enriched for genes that are down-regulated in post-mortem data (13). The relationship between the sign of MRI phenotype-related gene expression and the sign of post mortem transcriptional dysregulation in schizophrenia was non-random (sign test, P< 10^−12^).

Further investigation showed that the proteins coded by the PLS-genes formed a dense interaction network that was significantly enriched for a number of relevant GO biological processes and KEGG pathways. Proteins enriched for the GO term “adenylate cyclase-modulating G-protein coupled receptor (GPCR) signaling pathway” and for KEGG pathways “neuroactive ligand-receptor interaction” and “retrograde endocannabinoid signaling” were concentrated in a dense cluster of the PPI network. This GPCR signaling cluster includes multiple genes previously linked to pathogenesis and/or treatment of schizophrenia from diverse lines of evidence. Several of these genes have been implicated in anti-psychotic mechanisms of action, including DRD4 (18), HTR1 (34), NTSR1 (31) and ADRA2C (20). mRNA studies in post-mortem brain tissue also showed alterations in PTGER3, S1PR1, ITPR2, EDNRB and SSTR2 (13, 25, 27), and polymorphisms of DRD5, OPRM1 and CNR1 genes have been associated with susceptibility to schizophrenia (19, 23, 24).

The genetic support from GWAS and other data on sequence variation and from text mining was somewhat less compelling. Schizophrenia risk genes from the DISEASES dataset were enriched among genes that are normally overexpressed in frontal and temporal areas of reduced MS in psychosis; but the risk genes identified by the largest extant GWAS studies of schizophrenia were not significantly enriched among PLS-(or PLS+) genes. Nevertheless, the involvement of PLS-genes further down the causal pathway is still mechanistically revealing and potentially useful. Indeed, the remarkable density of therapeutically relevant genes in this small cluster suggests that surrounding genes may deserve further attention as novel anti-psychotic drug targets.

#### Methodological considerations

Some limitations of this study should be highlighted. The whole brain data on “normal” brain tissue expression of the genome were measured post mortem in 6 adult brains (mean age = 43 years) and not in age-matched subjects or patients with schizophrenia (such data are not currently available to our knowledge). Also, the transcriptional experiments we use to label genes as up- or down-regulated in schizophrenia were performed in regions of the parietal or prefrontal cortex (13), whereas the neuroimaging results are for the whole brain. We have used MRI data from 3 independent studies to measure MS networks but the studies used different scanning protocols, leading to estimation of morphometric similarity between regions on the basis of 7 MRI parameters that were measurable in all studies. Future work could usefully explore the opportunity to further improve sensitivity and reliability of the MS network biomarker of schizophrenia by optimising and standardising the MRI procedures to measure the most informative set of morphometric features.

## Methods

### Samples

We used MRI data from 3 prior case-control studies: the Maastricht GROUP study (35) from the Netherlands; the Dublin dataset which was acquired and scanned at the Trinity College Institute of Neuroscience as part of a Science Foundation Ireland-funded neuroimaging genetics study (“A structural and functional MRI investigation of genetics, cognition and emotion in schizophrenia”); and the publicly available Cobre dataset (36). The Maastricht and Dublin datasets were PSYSCAN legacy datasets. All patients satisfied DSM-IV diagnostic criteria for schizophrenia or other non-affective psychotic disorders. MRI data were quality controlled for motion artifacts, see SI. Demographic and clinical data on the evaluable samples are summarised in Table S1. The Euler number, which quantifies image quality (37), was not significantly different between groups in any of the studies but it was different between studies, indicating that the studies were ranked Dublin > Cobre > Maastricht in terms of image quality (Table S1). See SI for details.

### Morphometric similarity mapping

The T1-weighted MRI data (MPRAGE sequence) and the diffusion-weighted imaging (DWI) data from all participants were pre-processed using a previously defined computational pipeline (38). Briefly, we use the recon-all (39-42) and trac-all (43) commands from FreeSurfer (version 6.0). Following (5), the surfaces were then parcellated using an atlas with 308 cortical regions, derived from the Desikan-Killiany atlas (7, 44). For each region, we estimated 7 parameters from the MRI and DWI data: grey matter volume, surface area, cortical thickness, Gaussian curvature, mean curvature, fractional anisotropy (FA) and mean diffusivity (MD). Each parameter was normalised for sample mean and standard deviation before estimation of Pearson’s correlation for each pair of Z-scored morphometric feature vectors, which were compiled to form a 308 x 308 morphometric similarity matrix ℳ_*i*_ for each participant, *i = 1,…N* (5).

### Case-control analysis of MS networks

The global mean MS for each participant is the average of ℳ_*i*_. The regional mean *MS*_*i,j*_, for the ith participant at each region, *j* = 1,…, 308, is the average of the jth row (or column) of ℳ _*i*_. For global and regional MS statistics alike, we fit linear models to estimate case-control difference, with age, sex, and age x sex as covariates. P-values for case-control differences in regional MS were combined across all 3 studies, using Fisher’s method. The resulting P-value for each region which was thresholded for significance using the false discovery rate, *FDR <* 0.05, to control type 1 error over multiple (308) tests.

### Transcriptomic analysis

We used the AHBA transcriptomic dataset, with gene expression measurements in 6 postmortem adult brains (45) (http://human.brain-map.org), aged 24-57. Each tissue sample was assigned to an anatomical structure using the AHBA MRI data for each donor (46). Samples were pooled between bilaterally homologous cortical areas. Regional expression levels for each gene were compiled to form a 308 x 20, 647 regional transcription matrix (46). Since the AHBA only includes data for the right hemisphere for two subjects, in our analyses relating gene expression to MRI data we only consider the left hemisphere. We used PLS to relate the regional MS case-control differences (*t*-scores) to the post mortem gene expression measurements. PLS uses the gene expression measurements (the predictor variables) to predict the regional MS patient/control t-statistics from all 3 datasets (the response variables). The first PLS component is the linear combination of the weighted gene expression scores that has a cortical expression map that is most strongly correlated with the map of case-control MS differences.

We constructed PPI networks from the genes with PLS1 weights *Z <* –3 or *Z >* 3 using STRING (12) (our key results were robust to changing this threshold). We used DAVID (47, 48) to calculate enrichments of KEGG pathways and GO enrichments of biological processes for genes with *Z >* 3 or *Z <* –3, using a background gene list of 15,745 brain-expressed genes, see the SI (49).

We used a resampling procedure to test for enrichment of PLS-derived gene sets by genes previously associated with schizophrenia by transcriptional or sequence data (13) (14). The median rank of each risk gene set in the PLS gene list was compared to the median rank of 10,000 randomly selected brain-expressed gene sets (5).

Summary statistics from a meta-analysis of GWAS results from CLOZUK and PGC data were obtained from (15). To test PLS-derived genes for enrichment, we performed a gene set analysis on PLS+/PLS-using MAGMA v1.07b (50). First, gene-wide P values were calculated by combining the P-values of all SNPs inside genes, using a window of 35 kb upstream and 10 kb downstream of each gene to capture SNPs in approximate regulatory regions (51). We then performed one-tailed competitive gene set analysis and gene property analysis, using a background list of brain-expressed genes.

## ACKNOWLEDGEMENTS

This study was supported by grants from the European Commission (PSYSCAN - Translating neuroimaging findings from research into clinical practice; ID: 603196) and the NIHR Cambridge Biomedical Research Centre (Mental Health). The Co-bre data was downloaded from the Collaborative Informatics and Neuroimaging Suite Data Exchange tool (COINS; http://coins.mrn.org/dx) and data collection was performed at the Mind Research Network, and funded by a Center of Biomedical Research Excellence (COBRE) grant 5P20RR021938/P20GM103472 from the NIH to Dr. Vince Calhoun. SEM holds a Henslow Fellowship at Lucy Cavendish College, University of Cambridge, funded by the Cambridge Philosophical Society. PEV is a Fellow of MQ: Transforming Mental Health, grant number MQF17-24. KJW was funded by an Alan Turing Institute Research Fellowship under EPSRC Research grant TU/A/000017. The LTEXtemplate for this bioRxiv submission was created by Ricardo Henriques (and shared under Creative Commons CC BY 4.0).

