## Supplementary Information for "Cortical patterning of abnormal morphometric similarity in psychosis is associated with brain expression of schizophrenia related genes"

December 20, 2018

#### Contents

**1 Datasets** **2**

**2 Demographics** **2**

**3 Motion** **4**

**4 Global patient/control MS differences** **6**

**5 Regional patient/control MS differences in individual datasets** **6**

**6 Robustness of results** **8**

*PEV and ETB contributed equally to the work.

### 1 Datasets

Further details of the three datasets are given below. The datasets were chosen because they had both T1w MPRAGE and DWI images from 3T scanners for large (N > 100) sample sizes. In all three datasets, only subjects with both DWI and T1w images were included.

##### Maastricht GROUP

The Maastricht GROUP dataset comes from an MRI study in Maastricht, the Netherlands, led by the GROUP consortium. Patients were identified by screening caseloads of representative clinicians for inclusion criteria in selected representative geographic areas of the Netherlands and Belgium. The data was acquired using a 3T Siemens Magnetom Allegra head scanner. For more information, see [1]. 2 patients were excluded due to movement artifacts based on visual QC of the T1w data, leaving 83 patients and 68 control subjects. The diagnoses of the patients are heterogeneous and includes 57 patients with schizophrenia, 11 patients with psychotic disorder, 2 patients with brief psychotic disorder, 9 patients with schizoaffective disorder and 4 patients with schizophreniform disorder.

##### Dublin

The Dublin dataset was acquired and scanned in the Trinity College Institute of Neuroscience as part of a Science Foundation Ireland-funded neuroimaging genetics study ("A structural and functional MRI investigation of genetics, cognition and emotion in schizophrenia"). Patients were recruited through local clinical services whilst healthy control subjects reported no history of psychiatric disease. Both groups were recruited in the same geographical area through local advertisement and exclusion criteria for both groups included confirmed or suspected pregnancy, any history of neurological disorders or intellectual disability and substance misuse in the preceding 3 months. The data was acquired using a 3T Philips Intera Achieva scanner. 5 patients and 4 control subjects were excluded from the Dublin dataset due to movement artifacts based on visual QC of the T1w data. A large number of subjects also had to be excluded due to poor quality of DTI which led to failure to pass the pre-processing pipeline described below (51 control subjects and 9 patients). 82 control subjects and 33 patients remained, of whom 3 were diagnosed with schizoaffective disorder and 30 with schizophrenia.

##### Cobre

The Cobre dataset was obtained from the Neuroimaging Informatics Tools and Resources Clearinghouse (NITRC) website, and was provided by the Centers of Biomedical Research Excellence (COBRE; fcon_1000.projects.nitrc.orgindiretrocobre.html). In this dataset, a diagnosis of schizophrenia was made using the Structured Clinical Interview for DSM Disorders (SCID; Diagnostic and Statistical Manual of Mental Disorders, DSM-IV)(31). Exclusion criteria included confirmed or suspected pregnancy, any history of neurological disorders and a history of mental retardation. The data was acquired using a 3T Siemens scanner. 60 patients were diagnosed with schizophrenia and 9 with schizoaffective disorder.

### 2 Demographics

Table S1 gives the data demographics.

| | Maastricht GROUP | | Dublin | | Cobre | |
|---|---|---|---|---|---|---|
| | CON | PAT | CON | PAT | CON | PAT |
| Sample size | 68 | 83 | 82 | 33 | 77 | 69 |
| Age (years) | $29.4 \pm 10.3$ | $28.4 \pm 7.0$ | $33.5 \pm 12.6$ | $42.2 \pm 11.7$ | $38.0 \pm 12.0$ | $38.2 \pm 13.3$ |
| Sex (M) | 28 (41.2%) | 56 (67.5%) | 35 (42.7%) | 24 (72.7%) | 58 (75.3%) | 55 (77.5%) |
| PANSS total | $31.7 \pm 4.3$* | $44.3 \pm 13.8$* | N/A | N/A | N/A | $60.5 \pm 16.6$* |
| PANSS positive | $7.3 \pm 1.1$* | $9.9 \pm 4.1$* | N/A | N/A | N/A | $15.3 \pm 5.1$* |
| PANSS negative | $7.2 \pm 0.9$* | $11.1 \pm 5.6$* | N/A | N/A | N/A | $15.2 \pm 5.3$* |
| PANSS general | $17.3 \pm 2.7$* | $23.3 \pm 6.5$* | N/A | N/A | N/A | $30.1 \pm 9.9$* |
| SAPS (composite) | N/A | N/A | N/A | $12.9 \pm 16.4$* | N/A | N/A |
| SANS (composite) | N/A | N/A | N/A | $16.0 \pm 16.5$* | N/A | N/A |
| Euler number | $-95.7 \pm 49.0$ | $-96.5 \pm 53.9$ | $-35.5 \pm 18.7$ | $-30.9 \pm 19.6$ | $-66.6 \pm 36.4$ | $-66.0 \pm 31.0$ |

Table S1: Maastricht GROUP, Dublin and Cobre dataset demographics (after after subjects with poor data quality were excluded). *Dublin symptom scores were available for 24 patients only. Cobre symptom scores were available for 67 patients. Maastricht symptom scores were available for 62 control subjects and 65 patients.

### 3 Motion

To check for differences in motion and image quality between the patients and the control subjects we calculate the Euler number for each T1w image. This approach was proposed by [2] as a way to quantitatively assess image quality. Using a two-sided t-test we find no significantly significant differences in the Euler number between the two groups in any of the three datasets, as shown in Figure S1. We note that there are some differences in Euler number between datasets- for Maastricht the mean Euler number and standard deviation is $-96.2 \pm 48.9$, for Dublin we obtain $-34.3 \pm 18.9$ and for Cobre we obtain $-66.3 \pm 33.8$. Tentatively, this would suggest that the data quality is highest for the Dublin dataset and lowest for the Maastricht dataset, with the Cobre dataset in between the two. The exact extent to which the Euler number can be compared between datasets obtained using different scanners is unclear, for example [2] found that when predicting manually assessed image quality from the Euler number the most accurate classifications were obtained when classification threshold was allowed to vary by dataset, although accuracies of above 75% could still be obtained using a fixed threshold. Further work is needed in this area.

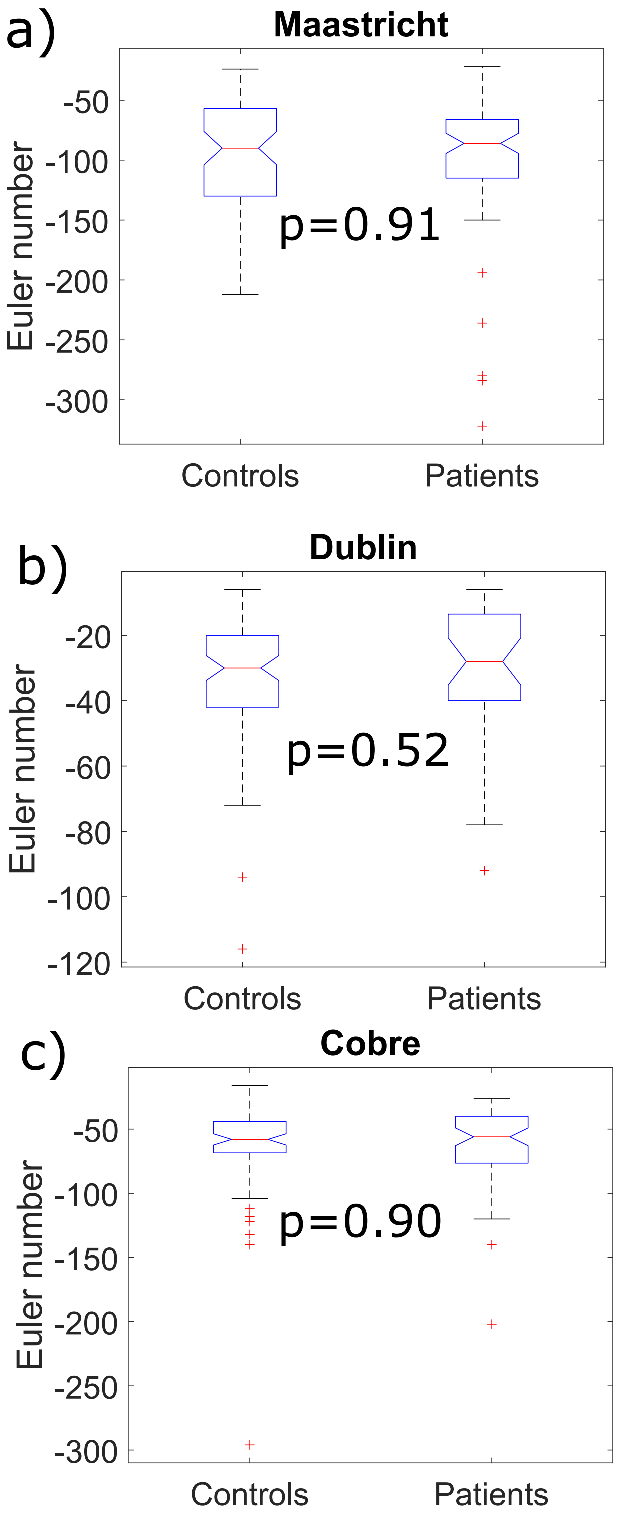

Figure S1: Box plots for the Euler number in the patient and control groups, for all three datasets.

### 4 Global patient/control MS differences

Figure S2 plots the global mean and distributions of regional morphometric similarity in all three datasets.

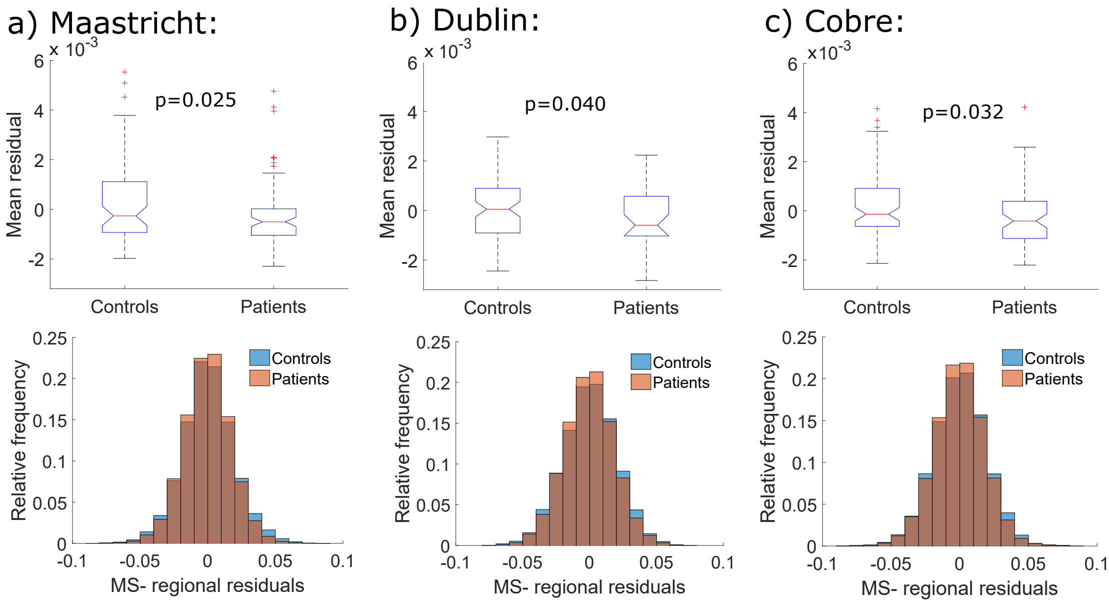

Figure S2: **Global mean and distributions of regional morphometric similarity.** Top panel: Box plots for global mean MS (after regressing sex and age) in controls and patients, in each of the three datasets (Maastricht, Dublin, COBRE). Bottom panel: Distributions of regional similarity strength, i.e., the average similarity of each region with all other regions, after regressing age and sex, for controls (blue) and patients (red), in each of the three datasets.

### 5 Regional patient/control MS differences in individual datasets

Figure S3 shows the patient/control MS t-statistics from each dataset, with regions where $P < 0.05$ only. Note that these results are shown before correcting for the false discovery rate, which is performed after combining the datasets. There are 90 significant regions in the Dublin dataset, 37 in the Cobre dataset and 30 in the Maastricht dataset, suggesting that the strongest signal is in the Dublin dataset.

In Figure S4, we plot the correlations between the t-statistics from the three different datasets, pairwise. As noted in the text, we find that the Dublin t-statistic is positively correlated with both the Maaastricht t-statistic (r=0.42, p< 0.001) and the Cobre t-statistic (r=0.47, p< 0.001), although the Maastricht and Cobre t-statistics are not correlated with each other (r=0.058, p=0.31).

After combining the datasets and correcting for FDR, 18 regions show statistically significant MS differences between control subjects and patients. The regions are listed in Table S2.

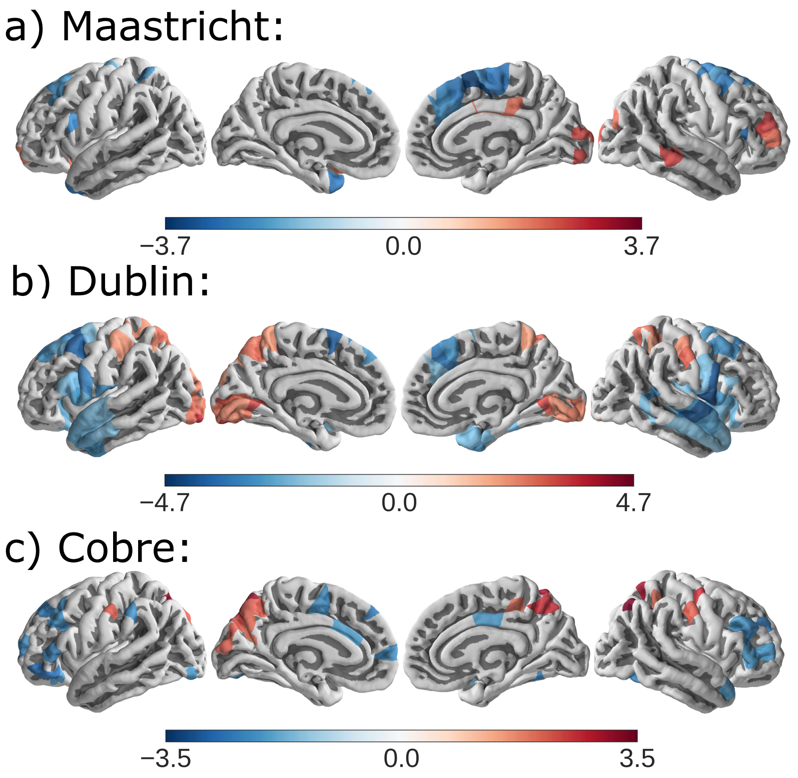

Figure S3: Patient/control MS t-statistics from a) Maastricht, b) Dublin and c) Cobre, with regions where $p < 0.05$ only, before FDR correction.

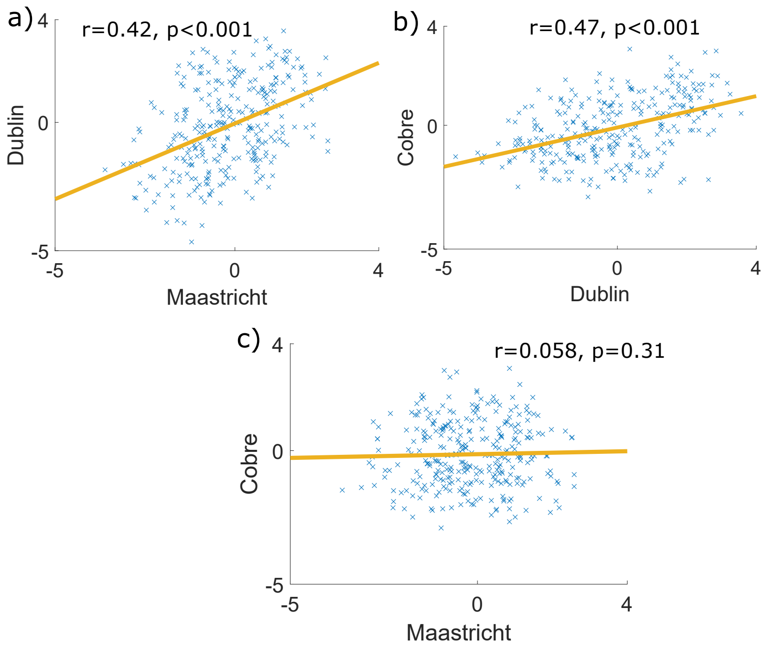

Figure S4: Correlations between the t-statistics from a) Dublin and Maastricht, b) Dublin and Cobre and c) Maastricht and Cobre.

| Name | x | y | z | Control MS | Patient MS | t-statistic | P-value |
|---|---|---|---|---|---|---|---|
| lh_caudalmiddlefrontal_part1 | -33.27 | 20.20 | 45.35 | 0.024 | 0.016 | -2.19 | 0.035 |
| lh_caudalmiddlefrontal_part4 | -39.72 | 11.34 | 48.85 | 0.022 | 0.013 | -2.27 | 0.015 |
| lh_precentral_part4 | -54.64 | 3.80 | 21.88 | 0.023 | 0.015 | -2.31 | 0.015 |
| lh_superiorfrontal_part8 | -10.01 | 10.76 | 61.43 | 0.012 | 0.002 | -2.27 | 0.015 |
| lh_superiorfrontal_part9 | -12.80 | 41.99 | 41.76 | 0.028 | 0.021 | -2.38 | 0.015 |
| lh_superiorfrontal_part11 | -17.72 | 30.84 | 48.03 | 0.021 | 0.011 | -2.02 | 0.049 |
| lh_superiorfrontal_part12 | -17.76 | 21.07 | 55.61 | 0.017 | 0.006 | -2.33 | 0.015 |
| lh_superiorparietal_part10 | -12.30 | -70.56 | 52.29 | -0.014 | -0.007 | 1.72 | 0.022 |
| rh_caudalmiddlefrontal_part4 | 36.44 | 12.22 | 48.85 | 0.016 | 0.007 | -2.43 | 0.015 |
| rh_parstriangularis_part2 | 43.06 | 25.40 | 5.94 | 0.001 | -0.008 | -2.11 | 0.035 |
| rh_postcentral_part6 | 54.62 | -14.84 | 36.20 | -0.018 | -0.011 | 2.01 | 0.049 |
| rh_precentral_part1 | 50.42 | 1.97 | 8.00 | 0.026 | 0.017 | -2.42 | 0.014 |
| rh_precentral_part3 | 54.26 | 5.64 | 24.20 | 0.019 | 0.010 | -2.45 | 0.015 |
| rh_superiorfrontal_part7 | 9.69 | 8.29 | 60.03 | 0.014 | 0.005 | -2.36 | 0.015 |
| rh_superiorfrontal_part11 | 9.20 | 24.39 | 53.69 | 0.020 | 0.012 | -2.18 | 0.019 |
| rh_superiorfrontal_part13 | 9.66 | 30.31 | 45.21 | 0.027 | 0.018 | -2.29 | 0.015 |
| rh_superiorparietal_part9 | 18.52 | -59.15 | 61.76 | -0.010 | -0.004 | 1.68 | 0.015 |
| rh_superiortemporal_part4 | 56.76 | -7.72 | -6.31 | 0.026 | 0.018 | -2.05 | 0.018 |

Table S2: Table giving details for the 18 statistically significant regions- anatomical labels, coordinates (in fsaverage, MNI305 space), mean MS value in control subjects and patients (averaged across datasets), mean t-statistic (averaged across datasets) and P-value after FDR correction.

### 6 Robustness of results

#### 6.1 Bootstrap within dataset

To assess the robustness of the regional MS results, we re-calculated the t-statistic for control/patient differences 1000 times with scrambled group labels. For each region we then calculate a P-value to assess whether the empirically calculated regional t-statistic is greater than the values from scrambled group labels (or less than in the case of a negative empirical t-statistic).

#### 6.2 Correlations between datasets

In order to assess the robustness of the correlations between datasets, we correlate the 1000 t-statistics obtained above (using scrambled group labels) pairwise between datasets. We then calculate a P-value based on the number of times the randomised correlations are greater than the real correlations observed between the datasets (a one-sided test). The results are given in Table S3. We find that the statistically significant correlations between the Maastricht and Dublin datasets and the Cobre and Dublin datasets reported in the main text are remarkably robust to this randomisation.

| Datasets | Pearson r | Pearson P-value | Bootstrap P-value |
|---|---|---|---|
| Maastricht/Dublin | 0.42 | $< 0.001$ | 0.0050 |
| Maastricht/Cobre | 0.058 | 0.31 | 0.36 |
| Dublin/Cobre | 0.47 | $< 0.001$ | $< 0.001$ |

Table S3: Pearson correlation values (r) for the correlations between t-statistics from the three datasets, alongside the corresponding Pearson correlation P-value and a P-value calculated using randomised group labels (see text for details).

#### 6.3 Spatial permutation

Finally, the correlation values reported above assume that the number of samples is equal to the number of regions, which is not the case because the number of regions is arbitrary (due to the resolution of the chosen parcellation) and non-independent (due to spatial autocorrelation amongst neighboring parcels). To address this issue, we use a regional spatial permutation test, as proposed by [3]. Here the idea is to compare the empirical correlation between the two t-statistics to null correlations generated by randomly rotating the spherical projection of one of the two spatial maps (as generated in FreeSurfer), before projecting it back on the brain surface. The rotated projection preserves both the spatial contiguity and the hemispheric symmetry of the empirical maps. See [3] for more details. The results are given in Table S4 and show that the correlations between the Maastricht GROUP and Dublin datasets and between the Dublin and Cobre datasets are robust to controlling for these effects.

| Datasets | Pearson r | Pearson p-value | Spatial permutation p-value |
|---|---|---|---|
| Maastricht/Dublin | 0.42 | $< 0.001$ | $< 0.001$ |
| Maastricht/Cobre | 0.058 | 0.31 | 0.38 |
| Dublin/Cobre | 0.47 | $< 0.001$ | $< 0.001$ |

Table S4: Pearson correlation values (r) for the correlations between t-statistics from the three datasets, alongside the corresponding Pearson correlation p-value and a p-value calculated using spatial permutation (see text for details).

##### 6.4 Outlier

We note that there is one subject which has outlier MD and FA values in the Cobre dataset. The data passed the pre-processing pipeline, however the output MD and FA values are substantially lower than the mean MD and FA values for the other subjects (more than 5 standard deviations), in all regions. This is the only subject for whom this is a problem, and there are no such subjects in either the Maastricht GROUP or Dublin datasets. If this subject is excluded from the analyses the results remain almost identical- the global morphometric similarity still decreases (p=0.034), as shown in Figure S5 b) and the t-statistics for the regional control/patient MS differences before and after excluding the subject correlate with Pearson correlation coefficient r=0.9973. This analysis confirms that the outlier subject does not drive our results and also suggests that our results are extremely robust to any individual subject.

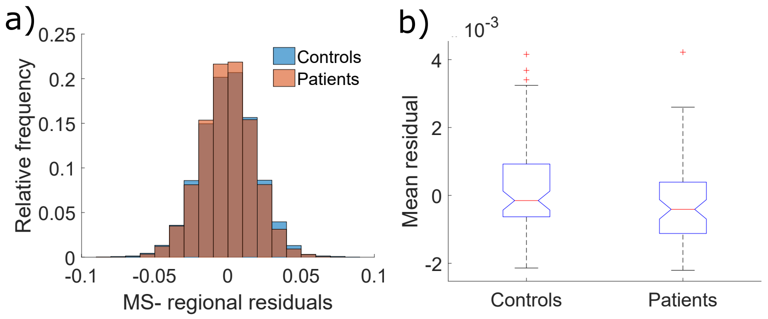

Figure S5: a) MS correlation distribution for Cobre with outlier subject excluded (see text for details). b) With the outlier subject excluded, mean MS is still reduced in patients compared to control subjects, as reported in the main text (p=0.034).

#### 7 Yeo networks and von Economo classes

To contextualise the regional MS case-control differences, we referred them to two prior classifications of cortical areas: the von Economo atlas of cortex classified by cytoarchitectonic criteria [4]; and the Yeo atlas of cortex classified according to resting state networks derived from function MRI [5, 3]. To do this we calculate t-statistics and corresponding P-values for regional MS differences averaged across all of the regions within a particular Yeo network or von Economo class. The results are shown in Tables S5 and S6. We find that across datasets there are reductions in MS in the ventral attention, frontoparietal and default mode Yeo networks, see Table S5. MS is also reduced in von Economo class 2 (association cortex).

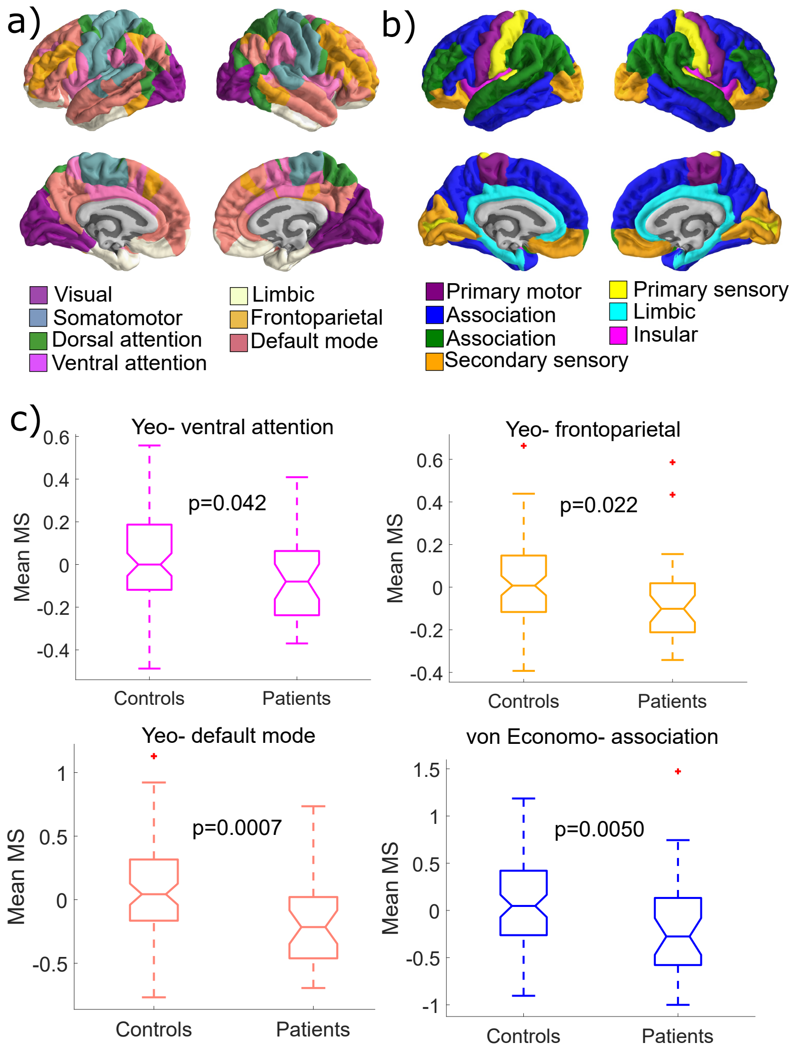

Figure S6: **Cytoarchitectonic and resting state network classification of case-control differences in regional morphometric similarity.** Brain plots coloured according to: a) the Yeo networks, b) the von Economo classes. c) Statistically significant control/patient differences in the Yeo networks and von Economo classes in the Dublin dataset. Specifically, we show the control/patient differences in Yeo network 4 (ventral attention), Yeo network 6 (frontoparietal), Yeo network 7 (default mode network) and von Economo class 2 (association cortex). The control/patient differences in these networks were also significant when results from all three datasets were combined- for details, see the text.

| Dataset | 1 (visual) | 2 (somatomotor) | 3 (dorsal attention) | 4 (ventral attention) | 5 (limbic) | 6 (frontoparietal) | 7 (default mode network) |
|---|---|---|---|---|---|---|---|
| Maastricht t | 1.1 | -1.7 | -1.3 | -1.9 | -0.20 | 0.89 | -1.0 |
| Maastricht p | 0.26 | 0.090 | 0.20 | 0.058 | 0.84 | 0.38 | 0.30 |
| Dublin t | 2.6 | 0.045 | 1.7 | -2.1 | -1.7 | -2.3 | -3.5 |
| Dublin p | **0.012** | 0.96 | 0.089 | **0.042** | 0.094 | **0.022** | **0.0007** |
| Cobre t | 0.69 | 1.1 | 1.5 | -2.0 | -0.66 | -3.2 | -2.2 |
| Cobre p | 0.50 | 0.29 | 0.15 | **0.043** | 0.51 | **0.0019** | **0.031** |
| Combined t | 1.5 | -0.20 | 0.63 | -2.0 | -0.85 | -1.5 | -2.2 |
| Combined p (after FDR) | 0.074 | 0.33 | 0.092 | **0.013** | 0.38 | **0.0041** | **0.0038** |

Table S5: t-statistics and p-values for regional MS control/patient differences averaged across each Yeo network. Results are shown for each dataset separately without FDR correction and then for the three datasets combined with FDR correction.

| Dataset | 1 (agranular, primary motor) | 2 (association) | 3 (association) | 4 (secondary sensory) | 5 (primary sensory) | 6 (limbic) | 7 (insular) |
|---|---|---|---|---|---|---|---|
| Maastricht t | -2.5 | -2.5 | 0.43 | 1.7 | 0.45 | -0.80 | -0.43 |
| Maastricht p | **0.013** | **0.012** | 0.67 | 0.10 | 0.65 | 0.42 | 0.67 |
| Dublin t | -1.1 | -2.9 | -1.2 | 2.5 | 2.0 | -0.072 | -1.7 |
| Dublin p | 0.28 | **0.0050** | 0.24 | **0.013** | **0.050** | 0.94 | 0.10 |
| Cobre t | 0.24 | -1.2 | -1.7 | -0.12 | 1.6 | -1.7 | -0.21 |
| Cobre p | 0.81 | 0.23 | 0.084 | 0.90 | 0.11 | 0.087 | 0.83 |
| Combined t | -1.1 | -2.2 | -0.83 | 1.4 | 1.3 | -0.86 | -0.77 |
| Combined p (after FDR) | 0.14 | **0.0072** | 0.27 | 0.12 | 0.14 | 0.41 | 0.45 |

Table S6: t-statistics and p-values for regional MS control/patient differences averaged across each von Economo class. Results are shown for each dataset separately without FDR correction and then for the three datasets combined with FDR correction.

### 8 Symptoms

#### 8.1 Convert Dublin dataset symptoms to PANSS

The Maastricht GROUP dataset and the Cobre dataset both report symptom information using the Positive and Negative Syndrome Scale for Schizophrenia (PANSS), whilst the Dublin dataset reports symptom information using the Scale for the Assessment of Positive Symptoms (SAPS) and the Scale for the Assessment of Negative Symptoms (SANS). [6] proposed formulae to convert between PANSS and SAPS/SANS measures:

$$\text{PANSS positive} = 11.1886 + (0.2587 \times \text{SAPS [Composite] Total score}) \quad \text{(S1)}$$

$$\text{PANSS negative} = 7.1196 + (0.3362 \times \text{SANS [Composite] Total score}) \quad \text{(S2)}$$

If these formulae are applied to the Dublin symptom scores shown in Table 1 of the main text, the Dublin PANSS positive score is $14.5 \pm 4.2$ and the Dublin PANSS negative score is $12.5 \pm 5.5$. Comparisons of symptom scores between datasets are given in Table S7. The results suggest that the Dublin and Cobre patients are more symptomatic than the patients from the Maastricht GROUP dataset in terms of positive symptom scores, whilst there is no statistically significant difference between the Dublin and Cobre datasets. The Cobre dataset is more symptomatic than either of the other two datasets in terms of negative symptom scores. The low symptom scores observed in the Maastricht dataset, suggesting that patients are, on average, not very symptomatic, is another possible reason that the Maastricht dataset $t$-statistic does not correlate with the Cobre dataset $t$-statistic, and the Maastricht dataset exhibits the fewest regions with significant control/patient differences in MS (in addition to the lower image quality of the Maastricht dataset).

| | PANSS positive | PANSS negative |
|---|---|---|
| Maastricht/Dublin | ✓(p< 0.001) | ✗(p=0.15) |
| Maastricht/Cobre | ✓(p< 0.001) | ✓(p< 0.001) |
| Dublin/Cobre | ✗(p=0.51) | ✓(p< 0.001) |

Table S7: Results from 2-sided t-tests to check for significant differences in the positive/negative symptom measures between the different datasets.

#### 9 Transcriptomic analysis

The PLS1 gene names and Z-score weights are provided in Dataset S1.

##### 9.1 Spatial permutation test- correlation between PLS1 and t-statistics

As above, we perform a spatial permutation test to assess whether the correlation between PLS1 and the t-statistics from the three datasets are robust to controlling for spatial autocorrelation amongst neighboring parcels. The results are given in Table S8. As reported in the main text, the correlation between PLS1 and the t-statistics from the Dublin and Cobre datasets are robust to this permutation test.

| Datasets | Pearson r | Pearson p-value | Spatial permutation p-value |
|---|---|---|---|
| Maastricht | 0.0060 | 0.94 | 0.55 |
| Dublin | 0.49 | $< 0.001$ | $< 0.001$ |
| Cobre | 0.37 | $< 0.001$ | 0.020 |

Table S8: Pearson correlation values (r) for the correlations between t-statistics from the three datasets, alongside the corresponding Pearson correlation p-value and a p-value calculated using spatial permutation (see text for details).

##### 9.2 PPI network analysis

We created PPI networks from the PLS- and PLS+ gene sets using the software STRING [7], with the highest confidence value of 0.9. We calculated GO enrichments for biological processes and KEGG pathway enrichments of the PLS- and PLS+ genes using the software DAVID [8, 9], with a background of 15745 brain-expressed genes. The background gene list is provided in Dataset S2 and was calculated by excluding probes which did not exceed the background noise in the AHBA dataset (intensity based filtering), as described by [10]. We used code from [10], with options.probeSelections = 'maxIntensity', inline with the maximum intensity approach used to derive our regional gene expression values [11].

##### 9.3 PPI network from genes with $Z < -3$

In the main text we showed GO enrichments for biological processes in the PPI network obtained from genes with $Z < -3$ (Figure 3). Two KEGG pathways are also enriched, as shown in Figure S7.

The full, high resolution PPI network from the genes with $Z < -3$, with the gene names labelled and coloured by GO enrichments for biological processes is given in Dataset S3. Significant GO enrichments for biological processes are listed in Dataset S1.

##### 9.4 PPI network from genes with $Z > 3$

In the main text we show that a PPI network from genes with $Z < -3$ has significantly more interactions than expected by chance and we also find significant enrichments in the network for certain KEGG pathways and GO enrichments of biological processes. A PPI network built from

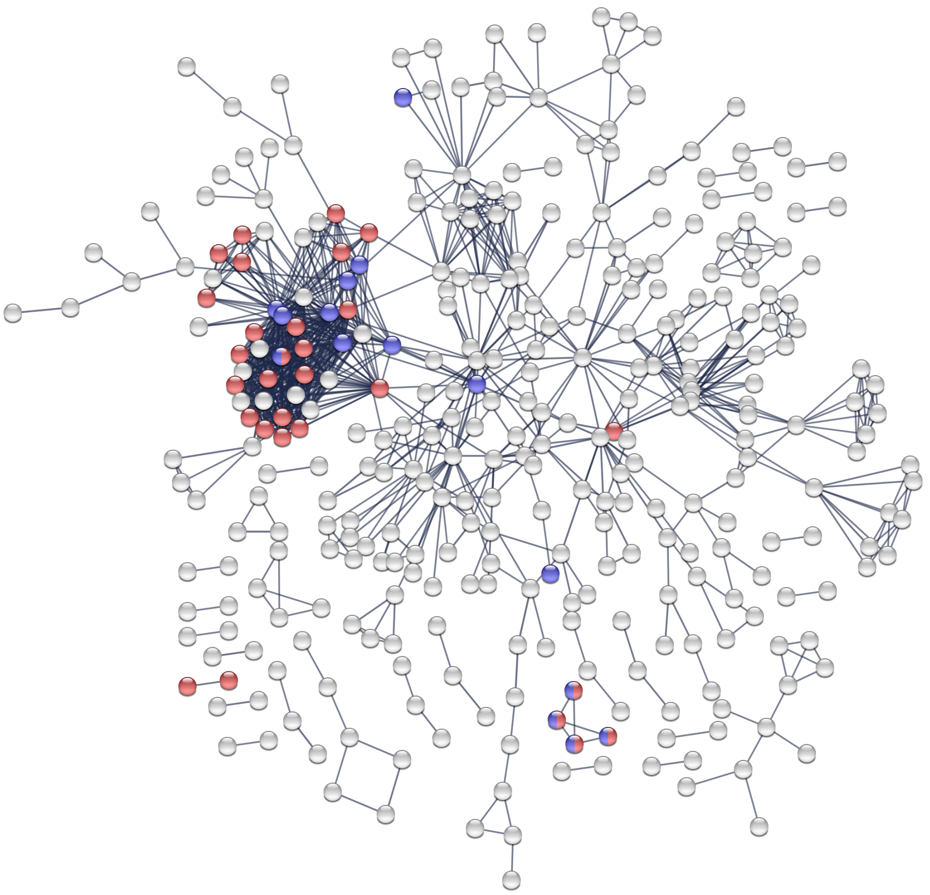

Figure S7: KEGG pathway enrichments, for PPI network from the PLS- gene set. Genes involved in the KEGG pathway 'neuroactive ligand-receptor interaction' are highlighted in red, genes involved in the KEGG pathway 'retrograde endocannabinoid signaling' are highlighted in blue.

genes with $Z > 3$ also has significantly more interactions than expected by chance. There are 1979 genes with $Z > 3$, 1802 of which are recognised by STRING. The resulting PPI network has 2808 edges, compared to an expected number of edges of 2542, giving a PPI enrichment p-value $< 1.0e-6$. The PPI network is shown in Figure S8, coloured by its GO enrichment for the biological process 'nucleic acid metabolic process'.

The full, high resolution PPI network from the genes with $Z > 3$, with the gene names labelled and coloured by GO enrichments for biological processes is given in Dataset S4.

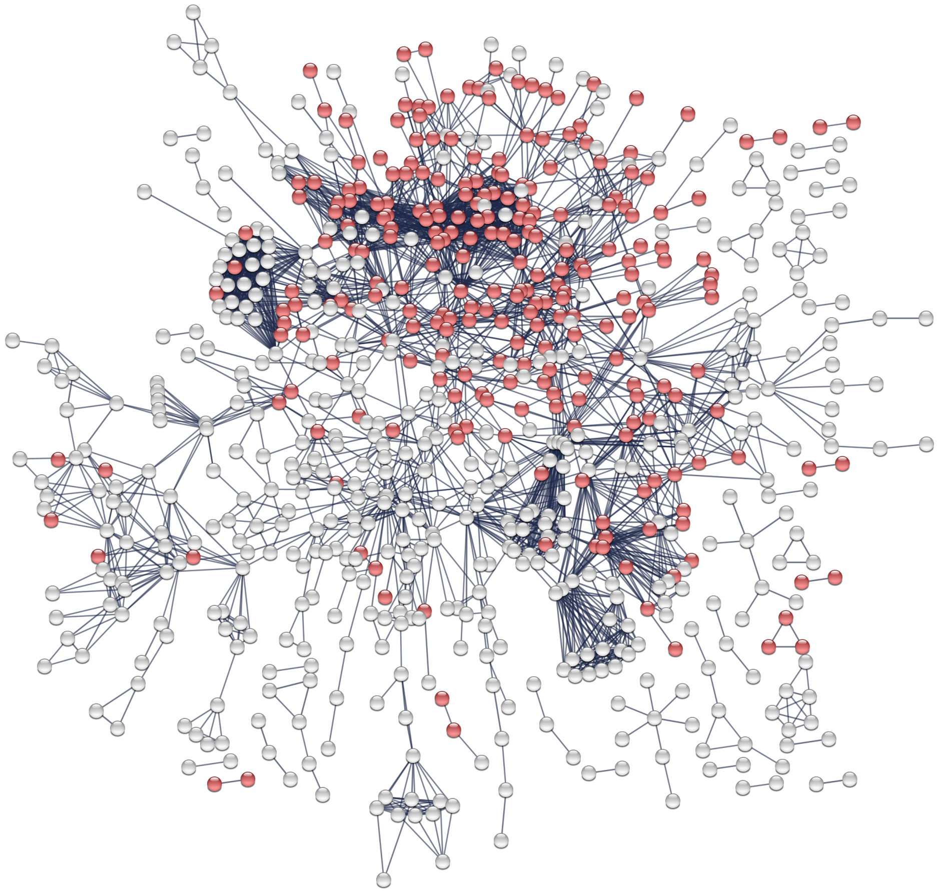

Figure S8: PPI network from PLS+ gene set. Genes highlighted in red are genes which belong to the enriched GO biological process- 'nucleic acid metabolic process'

#### 9.5 Specificity of Gandal gene enrichments

In the main text we report that in PLS1, genes which are over expressed in regions of increased and decreased MS in patients are enriched for the down and up regulated Gandal genes, respectively. Gandal et al [12] also includes lists of genes which are up and down regulated in the brain in a number of other disorders, namely bipolar disorder (BD), alcoholism (AAD), autism spectrum disorder (ASD) and major depressive disorder (MDD). Results for enrichments of these gene lists in PLS 1 are shown in Table S9. As in the main text, we use the background of brain-enriched genes. P-values are shown with no FDR correction. Our results show that the enrichment of genes which are up/down regulated in schizophrenia is not specific to abnormal gene expression in schizophrenia. Genes abnormally up regulated in autistic spectrum disorders (ASD), bipolar disorder and alcoholism were also significantly enriched among the genes that were positively or negatively weighted on PLS1, although genes abnormally up regulated in major depressive disorder were not. We note that there is substantial overlap between the lists of genes which are up/down regulated in the different psychiatric disorders, e.g. 20% of genes up-regulated in ASD were also up-regulated in schizophrenia.

| | Top of PLS1 | Bottom of PLS1 |
|---|---|---|
| AAD- up regulated | $< 0.0001$ | × |
| AAD- down regulated | × | 0.36 |
| ASD- up regulated | × | $< 0.0001$ |
| ASD- down regulated | $< 0.0001$ | × |
| BD- up regulated | × | 0.0013 |
| BD- down regulated | 0.27 | × |
| MDD- up regulated | × | 0.11 |
| MDD- down regulated | × | 0.30 |
| SZ- up regulated | × | $< 0.0001$ |
| SZ- down regulated | $< 0.0001$ | × |

Table S9: P-values for the enrichment of PLS 1 for lists of genes from several brain disorders obtained from the Gandal paper. P-values $> 0.5$ are not shown and P-values are not FDR corrected.

#### 9.6 Specificity of DISEASES gene enrichments

In the main text we report that in PLS1, genes which are over expressed in regions of decreased MS in patients are enriched for the DISEASES schizophrenia gene list (p=0.048). The DISEASES resource also includes lists of genes from other disorders, including Alzheimer's disease, ADHD, autistic disorder, bipolar disorder and mental depression, which allows us to check the specificity of our result. Results for enrichments of these gene lists in PLS 1 are shown in Table S10. We find that none of these gene lists is significantly enriched at either the top or the bottom of PLS 1 ($P > 0.05$, see the SI for details), apart from 'mental depression', hence our results show a high level of specificity to schizophrenia. The enrichment for 'mental depression' may be because patients with schizophrenia frequently also suffer from depression [13]. Methodological differences in diagnoses mean prevalence rates reported vary widely, but some reported rates are as high as 61% [14]. The symptoms and genetic risk factors between the two disorders are also known to overlap [13].

| | Top of PLS1 | Bottom of PLS1 |
|---|---|---|
| ADHD | × | 0.25 |
| Alzheimer's disease | × | 0.055 |
| Autistic disorder | × | 0.30 |
| Bipolar disorder | × | 0.27 |
| Mental depression | × | 0.015 |
| Schizophrenia | × | 0.048 |

Table S10: P-values for the enrichment of PLS 1 for lists of genes from several brain disorders obtained from the DISEASES resource. P-values $> 0.5$ are not shown and P-values are not FDR corrected.

#### 9.7 GAD

We also tested for significance of schizophrenia risk genes from the Genetic Association Database (GAD) [15, 16], which provides curated summary data on candidate gene and GWAS studies from published papers. To do this we use the same Method as for the Gandal and DISEASES gene lists. The GAD schizophrenia risk genes were downloaded from [16] were significantly enriched among genes negatively weighted on PLS1 (permutation test, $P = 0.0077$ before FDR correction), and not among genes positively weighted on PLS1 ($P > 0.05$).

Remarkably, a PPI network of the GAD schizophrenia genes shows a cluster of genes which is extremely similar to the cluster of genes in our PLS1 genes with $Z < -3$, as shown in Figure S9, adding confidence to the suggestion that this cluster of genes is highly relevant to schizophrenia. Figure S10 shows the GAD PPI network with the PLS1 genes with $Z < -3$ and $Z > 3$ highlighted. The genes with $Z < -3$ tend to be found in a different region of the GAD PPI network to the genes with $Z > 3$, suggesting that our imaging results differentiate between two sets of biological processes underlying schizophrenia.

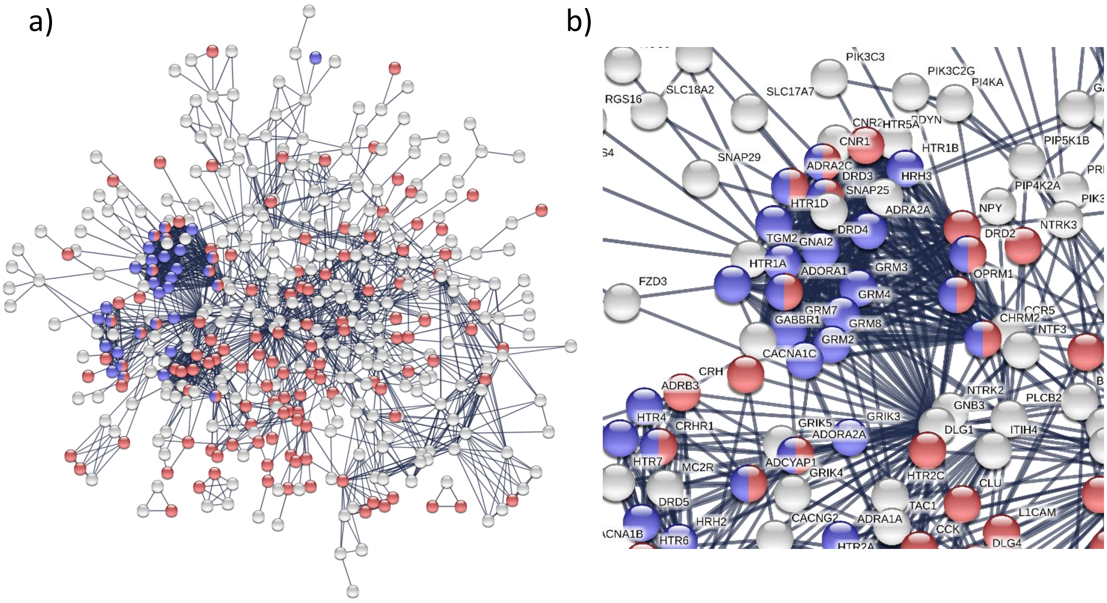

Figure S9: a) GAD PPI network, with GO enrichments for the terms 'nervous system development' and 'adenylate cyclase-modulating G-protein coupled receptor signaling pathway' highlighted in red and blue respectively. b) Zoom in on the cluster which closely resembles the cluster in the PPI network of PLS1 $Z < -3$ genes.

#### 9.8 FDR corrections

The P-values reported in the main text are given before FDR correction for multiple comparisons. Table S11 gives the P-values for the Gandal, DISEASES and GAD gene enrichments after FDR correction, which takes into account the fact that we have calculated enrichments at both the top and the bottom of each gene list (two comparisons). We note that the enrichments of the Gandal up-regulated and down-regulated genes at the bottom/top of PLS1, as well as the enrichment of the GAD genes at the bottom of PLS1 survive the FDR correction, although the enrichment of the DISEASES genes at the bottom of PLS1 does not. For the PGC+CLOZUK genes we performed three tests using the magma software [17]: gene set analyses of the PLS+ and PLS- genes as well as a gene property analysis which compared all of the (absolute) Z-scores from PLS1 with the P-values from the PGC+CLOZUK GWAS summary statistics. None of the P-values were significant either after or before FDR correction (before FDR correction, $P = 0.071$, $P = 0.94$, and $P = 0.93$ for gene set analysis for genes with $Z > 3$, $Z < -3$ and the gene property analysis, respectively).

#### 9.9 Overlap between lists of known schizophrenia genes

Table S12 gives details about the number of overlapping genes between the various gene lists we use. All gene lists overlap more than expected by chance. The strongest overlap is between the DISEASES and the GAD datasets (which may be because they come from overlapping sources). The number of genes expected to overlap is approximated as the number of genes in list 1 multiplied by the number of genes in list 2, divided by 20647 (the total number of genes in our analyses).

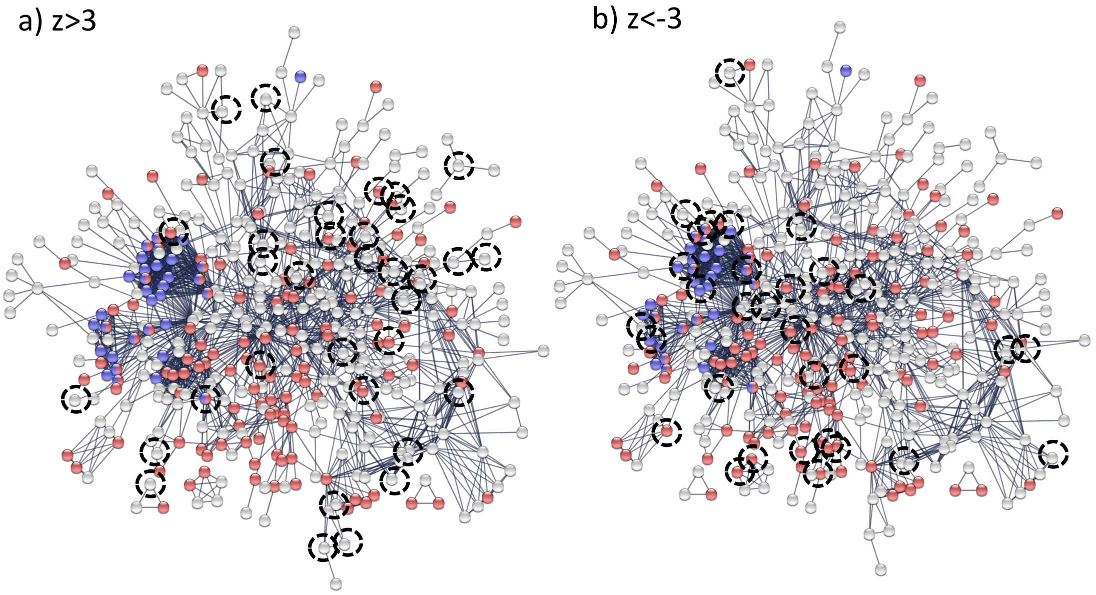

Figure S10: GAD PPI network, highlighting PLS1 genes with a) $Z > 3$ and b) $Z < -3$. GO enrichments for the terms 'nervous system development' and 'adenylate cyclase-modulating G-protein coupled receptor signaling pathway' are highlighted in red and blue respectively.

| | Top of PLS1 | Bottom of PLS1 |
|---|---|---|
| Up regulated genes from Gandal et al [12] | × | P< 0.001 |
| Down regulated genes from Gandal et al [12] | P< 0.001 | × |
| DISEASES | × | × |
| GAD | × | P=0.015 |

Table S11: Summary of the P-values related to the enrichment of the gene lists in PLS1 and PLS1 reversed, after FDR correction. P-values > 0.05 after FDR correction are not shown to aid readability.

| Gene lists | No. overlap genes | No. genes expected to overlap | No. overlap genes/expected overlap |
|---|---|---|---|
| Gandal up-regulated (849 genes) and GAD (674 genes) | 48 | 28 | 1.7 |
| Gandal down-regulated (1194 genes) and GAD (674 genes) | 69 | 39 | 1.8 |
| Gandal up-regulated (849 genes) and DISEASES (130 genes) | 9 | 5 | 1.7 |
| Gandal down-regulated (1194 genes) and DISEASES (130 genes) | 12 | 8 | 1.6 |
| DISEASES (130 genes) and GAD (647 genes) | 50 | 4 | 12 |

Table S12: Table with details about overlapping genes in the lists of known schizophrenia genes.

##### 9.10 Extended discussion of GPCR gene cluster

The PPI network from the genes which are over-expressed in regions of decreased MS is enriched for a number of relevant GO biological processes and KEGG pathways. Interestingly, the enrichments for the GO term "adenylate cyclase-modulating G-protein coupled receptor signaling pathway" and both significantly enriched KEGG pathways "neuroactive ligand-receptor interaction" and "retrograde endocannabinoid signaling" were concentrated in a tight cluster of the PPI network. Here we extend our discussion of this gene cluster. We note that the cluster includes multiple genes previously linked to schizophrenia from diverse lines of evidence across a number of genotypic and phenotypic levels. For example, polymorphisms of the DRD5, OPRM1 and CNR1 genes have each been associated with susceptibility to schizophrenia [18, 19, 20]. Microarray screening and real-time PCR of lymphocyte gene expression identified decreasing NPY1R and GNAO1 in individuals with schizophrenia compared to unaffected family controls [21]. Centrally, mRNA studies in post-mortem brain tissue showed alterations in PTGER3, S1PR1, ITPR2, EDNRB, SSTR1 AND SSTR2 [22, 23, 12]. Several genes in this cluster have also been implicated in therapeutic approaches to schizophrenia, including DRD4 which codes for the dopamine receptor 4 and is a target for drugs that treat schizophrenia and Parkinson's disease. HTR1 codes for the serotonin 1A receptor (or 5-HT1A receptor) and the 5-HT1A receptor partial agonist properties of a number of atypical antipsychotics have been shown to enhance their clinical efficacy [24]. NTSR1 is a high-affinity receptor for neurotensin, which has been shown to selectively modulate dopaminergic neurotransmission and which was found to be reduced in the CSF and post-mortem brain tissue in schizophrenia [25, 26]. Central administration of neurotensin was also observed to produce effects similar to those of atypical antipsychotics [27]. Finally, ADRA2C is a candidate gene for schziophrenia because it binds clozapine, an atypical antipsychotic medication widely prescribed for treatment-resistant schizophrenia [28]. We note that many of the above genes were not identified in most recent GWAS studies and therefore may not be implicated at the level of polymorphisms. Nevertheless, their involvement further down the causal pathway, at the epigenetic level is still mechanistically revealing and potentially useful in practice. Indeed, the remarkable density of therapeutically relevant genes in this small cluster suggest that surrounding genes may deserve further attention.
